# Acute ethanol intoxication induces preferred loss of presynaptic boutons devoid of mitochondria *in vivo*

**DOI:** 10.1101/536334

**Authors:** Jil Protzmann, Astha Jaiswal, Karl Rohr, Thomas Kuner, Sidney Cambridge, Johannes Knabbe

## Abstract

Acute alcohol intoxication is frequently observed in modern societies and carries a vast burden, ranging from traffic accidents to transient memory loss. Despite years of intense research, the effects of acute ethanol intoxication on brain function remain incompletely understood. Here, we studied the effect of acute ethanol intoxication on axonal organelle trafficking and presynaptic structure using *in vivo* two photon microscopy in anesthetized mice. After a single intraperitoneal injection of ethanol, inducing a blood alcohol concentration of roughly 250 mg/dl, the axonal mitochondrial mobility was doubled while dense core vesicle mobility remained unaffected. Simultaneously imaging mitochondria and presynaptic boutons revealed that unoccupied presynaptic boutons perished more frequently after ethanol exposure, while boutons stably occupied with mitochondria mostly persisted. Our results define a novel mechanism of ethanol action and may explain difficulties in permanently storing new memories after episodes of intense ethanol consumption with a loss of synapses.

## Introduction

Research in the last years identified numerous direct molecular targets of ethanol in the brain, including ligand-gated ion channels (e.g. NMDAR (Kuner, Schoepfer and Korpi 1993, Lovinger, White and Weight 1989); GABA_A_R (Marszalec et al. 1994); GlyR (Soderpalm, Lido and Ericson 2017); nAChR (Cardoso et al. 1999); others (Dopico and Lovinger 2009)), potassium channels (e.g. BK channels (Dopico, Bukiya and Martin 2014); GIRK channels (Bodhinathan and Slesinger 2013)), sodium channels (Horishita and Harris 2008), enzymes (e.g. ADH, (Goto et al. 2015)) and signaling factors (e.g. cAMP (Rabin and Molinoff 1981)). Ethanol differentially modulates these targets and thereby generates a multitude of biological effects (for review see Lovinger and Roberto (2013) or Abrahao, Salinas and Lovinger (2017)). Therefore, understanding ethanol-induced behavioral changes in mice or humans on the basis of these mechanisms of ethanol action remains challenging. A multiplicity of ethanol actions, rather than a single or a few distinct mechanisms, may underlie the behavioral effects of ethanol intoxication.

An effect of ethanol administration on organelle transport in neurons has been shown in the vagus nerve (Laduron and De Witte 1987), sciatic nerve (McLane (1990); Malatova and Cizkova (2002)) and sural nerve (McLane et al. 1992) of chronic ethanol-fed rats, though some of the results are inconsistent. Only recently, Iacobucci and Gunawardena (2018) could show an increase in axonal movement of atrial natriuretic factor-containing dense core vesicles (DCV) in motor neurons of *Drosophila* larvae caused during acute ethanol intoxication. DCVs, like mitochondria, are transported on microtubule tracks with motor proteins of the kinesin and dynein superfamilies (Zahn et al. (2004); Hollenbeck (1996)). Regulation of transport is activity dependent for both DCV and mitochondria. Elevated levels of intracellular calcium ([Ca^2+^]_i_) induces phosphorylation of synaptotagmin in the DCV membrane and thus immobilization of DCVs in presynaptic boutons (Bharat et al. 2017). Similarly, increased [Ca^2+^]_i_ at active zones of presynaptic terminal leads to the detachment of mitochondria from microtubules (Yi, Weaver and Hajnoczky (2004); Fransson, Ruusala and Aspenstrom (2006); Macaskill et al. (2009)) as well as to the binding of the anchor protein syntaphilin (Chen and Sheng 2013). These processes produce a pool of stationary mitochondria positioned at sites of constantly high energy demand, while the pool of mobile mitochondria allows for a flexible adaptation to changing metabolic needs (Hollenbeck and Saxton 2005). Two essential roles of mitochondria in neurons are rapid Ca^2+^ sequestration (Billups and Forsythe 2002) as well as ATP production (Rich and Marechal 2010). Interfering with mitochondrial mobility in axons abolished synaptic transmission during trains of action potentials (Verstreken et al. 2005). Exogenous ATP could rescue synaptic transmission, suggesting that mitochondrial ATP supply is crucial for sustained synaptic transmission under intense stimulation. However, only approximately half of all presynaptic boutons were shown to contain mitochondria (Chang, Honick and Reynolds 2006, Kang et al. 2008, Smith et al. 2016, Smit-Rigter et al. 2016). Smit-Rigter et al. (2016) observed that those boutons containing mitochondria were significantly more likely to be stable over a time period of 4 days. Qiao et al. (2016) analyzed the long-term stability of presynaptic boutons and could show that newly formed synaptic boutons perished more often in the subsequent 2-3 weeks than pre-existing synaptic boutons. Additionally, 80 % of axonal boutons survived 12 months arguing for subsets of presynaptic boutons that are either frequently turned over or persist for months to years.

In this study we aim to characterize mitochondrial and DCV transport as well as structural plasticity in mice *in vivo* in response to acute ethanol intoxication. We found that ethanol doubles mitochondrial mobility and triggers the loss of presynaptic boutons not occupied by mitochondria. These findings may provide new explanations for the effect of acute ethanol intake on learning and memory.

## Methods

### Ethical approval

Animal experiments were carried out in accordance with the guidelines of the European Science Foundation 2001 and were approved by the Governmental Supervisory Panel on Animal Experiments of Baden Wuerttemberg in Karlsruhe. Utmost priority was to avoid or minimize animal suffering, promote animal welfare and reduce the number of animals used. 8-week old male, wild-type (wt) C57BL/6N mice (Charles River Laboratories) were housed singly in an individually-ventilated cage system (ZOONLAB) at a 12 h dark/light cycle. The animal room was tempered to 23 °C with a relative humidity of 55 %. The mice had access to water and chow *ad libitum*, except for imaging sessions.

### Genetic labelling of mitochondria, DCVs and presynaptic boutons

In house produced (Schwenger and Kuner 2010) recombinant adeno-associated viruses (rAAVs) of the chimeric 1/2 serotype were used in order to label axonal organelles and presynaptic boutons in neurons *in vivo*. Mitochondria were labelled with GFP fused to the first 35 amino acids of the mitochondria-targeting sequence of cytochrome *c* oxidase subunit VIII leading to enrichment of GFP in the mitochondrial matrix (Plasmid was kindly provided by Dr. Ken Nakamura, see Berthet et al. 2014). Mitochondria-targeted GFP (mitoGFP) was expressed under the control of the EF-1 alpha promoter. As the construct contained a Cre-inducible double-floxed inverse open reading frame (DIO), Cre-recombinase was necessary for expression of the insert (EF-1α-DIO-mitoGFP). rAAVs encoding for Cre-recombinase under the control of the synapsin promoter (Syn-Cre) were always co-injected leading to a restricted expression of mitoGFP in neurons. For the labelling of dense-core vesicles (DCV), the DCV cargo protein neuropeptide-Y (NPY) fused to the fluorescent protein venus was used under the control of the synapsin promoter. The plasmid for Syn-NPY-venus was kindly provided by Dr. Matthijs Verhage. DCV labelling was combined with axonal labelling by expressing mCherry under the CAG promoter (Syn-mCherry). Labelling of presynaptic boutons was achieved by expressing synaptophysin (SyPhy) fused to mCherry under the control of the CAG promoter (CAG-SyPhy-mCherry).

### Craniectomy, viral injection and chronic cranial window implantation

The protocol was adapted from Holtmaat et al. (2009) as well as Boffi et al. (2018). Mice were deeply anaesthetized by intraperitoneally (i.p.) injecting a mixture of 0.48 µl fentanyl (1 mg/ml; Janssen), 1.91 µl midazolam (5 mg/ml; Hameln) and 0.74 µl medetomidin (1 mg/ml; Pfizer) per gram body weight each and mounted in a stereotaxic apparatus (EM70G, David Kopf Instruments). Mice were kept on 35°C by a feedback-controlled heating pad throughout the surgery. The eyes were covered with eye ointment (Bepanthen with 5% Dexpanthetol, Bayer Vital GmbH) to prevent drying-out. Standard aseptic procedures were followed during all surgeries. The local anesthetic Xylocain (1%, AstraZeneca) was injected subcutaneously (s.c.) at the site of surgery and the cranium was exposed by removing a flap of the scalp, approximately 1 cm², as well as the subjacent gelatinous periosteum. Levelling of the mouse skull to a coordinate system centered at bregma was performed by eye using a monocle and a digital indication (both David Kopf Instruments). A small hole was drilled into the skull with a dental drill (drill: EXL-40, Osada Inc.; boring head: 1104005, Komet Dental) and the right-hemisphere medio-dorsal nucleus of the thalamus (MD thalamus) was targeted using the coordinates 0.82 AP, 1.13 ML and 3.28 DV (Paxinos and Franklin 2013) mm relative to bregma. Approximately 800 nl of a mixture of rAAVs (1:1:1 mitoGFP, Cre, SyPhy-mCherry OR 1:1 NPY-venus, mCherry) was slowly injected. Directly after the viral injection, a larger craniectomy was performed. Therefore, a circular part of the skull, with a diameter of 6.5 −7 mm and centered above the injection site, was removed by drilling a deep groove into the outline of the circle. The dura mater was carefully removed with a very fine forceps (straight Dumont #5, Fine Science Tools Inc.). A glass coverslip (6 mm diameter, #0) was placed inside the opening and was, together with a 3D-printed round plastic holder, cemented to the scull will dental acrylic cement (glue: Cyano, Hager Werken; powder: Paladur, Heraeus). Mice received i.p. a mixture of 1.86 µl naloxon (0.4 mg/ml; Inresa), 0.31 µl flumazenil (0.1 mg/ml; Fresenius Kabi), 0.31 µl antipamezole (5 mg/ml; Pfizer) in 3.72 µl Saline (0.9 %; Braun) each per gram body weight to antagonize the anesthesia. For pain treatment mice received an s.c. injection of 150 µl Carprofen (50 mg/ml; Bayer Vital GmbH) diluted in saline every 12 h for the next two days. Mice rested for 5-8 weeks to ensure high rAAV expression and an eased inflammatory glial reaction (Holtmaat et al. 2009).

### In vivo two photon microscopy of anesthetized mice and ethanol injection

Two photon imaging (Denk et al. 1994) was conducted as described in Knabbe et al. (2018) using an upright TriM Scope II microscope (LaVision BioTec GmbH) equipped with a pulsed Ti:Sapphire laser (Chameleon Ultra II; Coherent). A wavelength of 960 nm was used to excite mitoGFP OR simultaneously NPY-venus and cytosolic mCherry. Imaging of mitoGFP with 960 nm and SyPhy-mCherry with 1050 nm was serially done. The emitted signals were separated using a 560 nm dichroic mirror (Chroma, Bellows Falls) and appropriate filter sets (GFP, venus: 535/70 nm emission filter (Chroma, Bellows Falls); mCherry: 645/75 nm emission filter (Semrock, Rochester)). Imaging was done using a 25x water immersion objective (1.1 NA, Nikon) and the emitted fluorescence was collected with photo multiplier tubes (PMTs, H7422-40-LV 5M; Hamamatsu). Mice were anesthetized in an acrylic box with 5 % isoflurane (Henry Schein) in medical O_2_ (Air Liquid Medical GmbH) using a vaporizer (Vapor 2000, Dräger). The animal was head-fixed into a custom-built holder screwed onto the intravital microscope stage. Vessels were used for orientation and relocating to the same imaging site during every imaging session. During the imaging session, isoflurane was maintained at 0.8 – 1.5%. Breathing rate was under constant monitoring using an infra-red camera (ELP 1080P 2.0 Megapixel USB camera, Ailipu Technology Co.) and an infra-red LED light source (48 LED Illuminator, Sonline). Body temperature of the animal was kept at 37 °C by a heating pad (ExoTerra, HAGEN Deutschland GmbH).

Layer 1 of motor cortex, 50-100 µm away from the pial surface was imaged. ImSpector (LaVision BioTec GmbH) was used for microscope control and image acquisition. When imaging mitochondria, frames were taken from an area covering 436 x 436 µm, at 1024 x 1024 pixel resolution (0.426 µm/pixel resolution) with a frame rate of 0.61 Hz using a galvanometer scanner. During acquisition of mitochondrial transport the focal plane was rapidly alternated using a piezo-motor (LaVision BioTec GmbH) allowing the acquisition of three focal planes at each time point resulting in an effective frame rate of 0.2 Hz per image plane. When imaging DCVs, frames were taken from an area covering 200 x 200 µm, at 1071 x 1071 pixel resolution (0.19 µm/pixel resolution) with a frame rate of 0.94 Hz using the galvanometer scanner. Each acquired time-lapse of mitochondrial transport was of 10 min duration and 5 min of DCV transport. Baseline time-lapses were recorded at three time points before i.p. injection of 500 µl of saline or 20 % ethanol in water. Time-lapses up to 4 h post-injection were acquired. The ethanol injection induced blood alcohol concentrations of approximately 250 mg/dl (Eisenhardt et al. 2015). Three additional time-lapses were acquired 24 h post-ethanol injection. 3D-stacks covering 436 x 436 x 50 µm, at 2731 x 2730 pixel resolution (0.16 µm/pixel resolution) and 1 µm z-steps were acquired pre, 4 h and 24 h post saline and ethanol injection in order to dissolve structural changes of mitochondria and presynaptic boutons.

### Immunohistochemistry

Mice were transcardially perfused with 4 % paraformaldehyde (Sigma Aldrich) and brains were carefully removed from the skull. For immunohistochemical stainings 80 µm coronal sections were cut with a microtome (Leica VT100S, Leica). Therefore, brain sections were incubated in PBS containing 5 % natural goat serum (NGS) and 1 % Tritum-X100 for 2 h, before incubating the slices in PBS containing 1 % NGS, 0.2 % Triton-X100 and the primary antibody over night at 4 °C. Slices were washed in PBS containing 2 % NGS. Secondary antibodies were incubated for 2-3 h at room temperature. Used primary antibodies: α-Translocase of the outer mitochondrial membrane (TOM20, 1:500; Santa Cruz), α-Synaptophysin 1 (SyPhy1, 1:500; Synaptic Systems), α-Chromogranin A (Chr-A, 1:500; Synaptic Systems). Alexa Fluor antibodies (1:500; Invitrogen) were used as secondary antibodies. Slices were mounted with Mowiol (SigmaAldrich) and imaged on a Leica DM6000 wide-field epifluorescence light microscope with 10x/0.4 (magnification/NA) or 63x/1.30 glycerol-immersion objectives or on a Leica SP8 scanning confocal microscope using a 63x/1.40 oil-immersion objective.

### Data processing and analysis

For image processing and analysis, the open-source image analysis software Fiji was used. Time-lapse images were registered in *xy* using the Fiji plugin “moco” (Dubbs, Guevara and Yuste 2016). Average intensity projections of time-lapse images were used to display stationary cell organelles as well as the outlines of axons, while maximum intensity projection was used to identify axons displaying active cell organelle trafficking. For mitochondrial transport, only active axonal stretches (ROIs) for analysis were selected. A total of 15 ROIs was analyzed per focal plane. Kymographs were generated with the Multi Kymograph tool (line width = 3) in Fiji. Kymographs revealed the number of stationary and mobile mitochondria, from which a mobility ratio was calculated. Kymographs were used to calculate the mean velocity of a mitochondrial track by using a Fiji macro written by Alessandro Moro. DCV transport was automatically tracked (see next section) and analyzed using self-written routines in Matlab (MathWorks). For the mobility analysis, DCV tracks were defined as mobile, if the total movement distance was above or equal to 5µm and stable if the movement distance was below 5µm. Mobility was calculated as mobile-fraction / (stable fraction + mobile fraction). Mean velocity was calculated on the mobile fraction only.

The mitoGFP and SyPhy-mCherry 3D-stacks were registered by manual selection of corresponding points and calculation of the resulting transformation in Matlab. For analyzing the structural plasticity of mitochondria and presynaptic boutons axons clearly visible in all time points (pre, 4 h and 24 h post) were chosen. Signals 1.5 times higher than the background of the axon were counted as mitochondria or presynaptic boutons, respectively.

### Automatic tracking of dense core vesicles

To quantify the mobility of vesicles in the mouse brain data, we used an automatic particle tracking approach, which performs multi-frame association by exploiting information from multiple time points (Jaiswal et al. 2016). In each image frame of a video, vesicles were detected by applying the spot-enhancing filter, which is based on the Laplacian of Gaussian operator (Sage et al. 2005). We used two different size thresholds yielding two sets of detections for small and large size spots at each time point of an image sequence. Detected spots of small and large size were used for tracking, while for initialization of trajectories only large spots were used. A Kalman filter (Kalman 1960) was employed to determine predictions about the state of vesicles using a two motion model comprising random walk and directed motion. For finding associations between the predictions and the detections at subsequent time points, a multi-frame approach was used, which is based on a graph theoretical formulation and supports one-to-one associations, many-to-one, and one-to-many associations. The optimal associations were computed by solving a linear program (Jaiswal et al. 2016). The state of vesicles was computed by the Kalman filter based on the resulting associations and the selected motion model. The same parameter setting was used for all image sequences of a mouse dataset. The parameter settings for datasets of different mice were slightly adapted. Short trajectories were excluded from the statistical analysis (we used a threshold of 5 frames). Broken trajectories were automatically merged on the basis of their spatial and temporal distances (we used thresholds of 15 pixels and 15 frames, respectively).

### Statistical analysis

Statistical analysis was performed in Prism (GraphPad Software). For parametric tests, normal distribution was tested using D’Agostino & Pearson normality test. For analysis either goodness of fit, linear regression, Mann-Whitney test, one-way ANOVA, two-way repeated measures ANOVA or Kruskal-Wallis test were performed. For avoiding α-error accumulation, results were adjusted using post-hoc Sidak’s multiple comparisons test (two way repeated-measures ANOVAs), post-hoc Dunn’s multiple comparisons test (Kruskal-Wallis test) or post-hoc Dunnett’s multiple comparisons test (one way-ANOVA). F values as well as degrees of freedom (DF; F (DFn, DFd)) are provided for ANOVAs. Statistical significance was assumed for *p* < 0.05. All data is displayed as mean ± standard deviation (SD) if not indicated otherwise (e.g. as mean ± standard error of the mean (SEM)).

## Results

### Experimental design

In the present study, we investigated the effect of acute ethanol intoxication on axonal organelle transport of mitochondria as well as DCVs and alterations in turnover of presynaptic boutons *in vivo*. For two-photon imaging of the thalamic projections, a chronic cranial window was implanted (Figure 1A).

**Figure 1:**
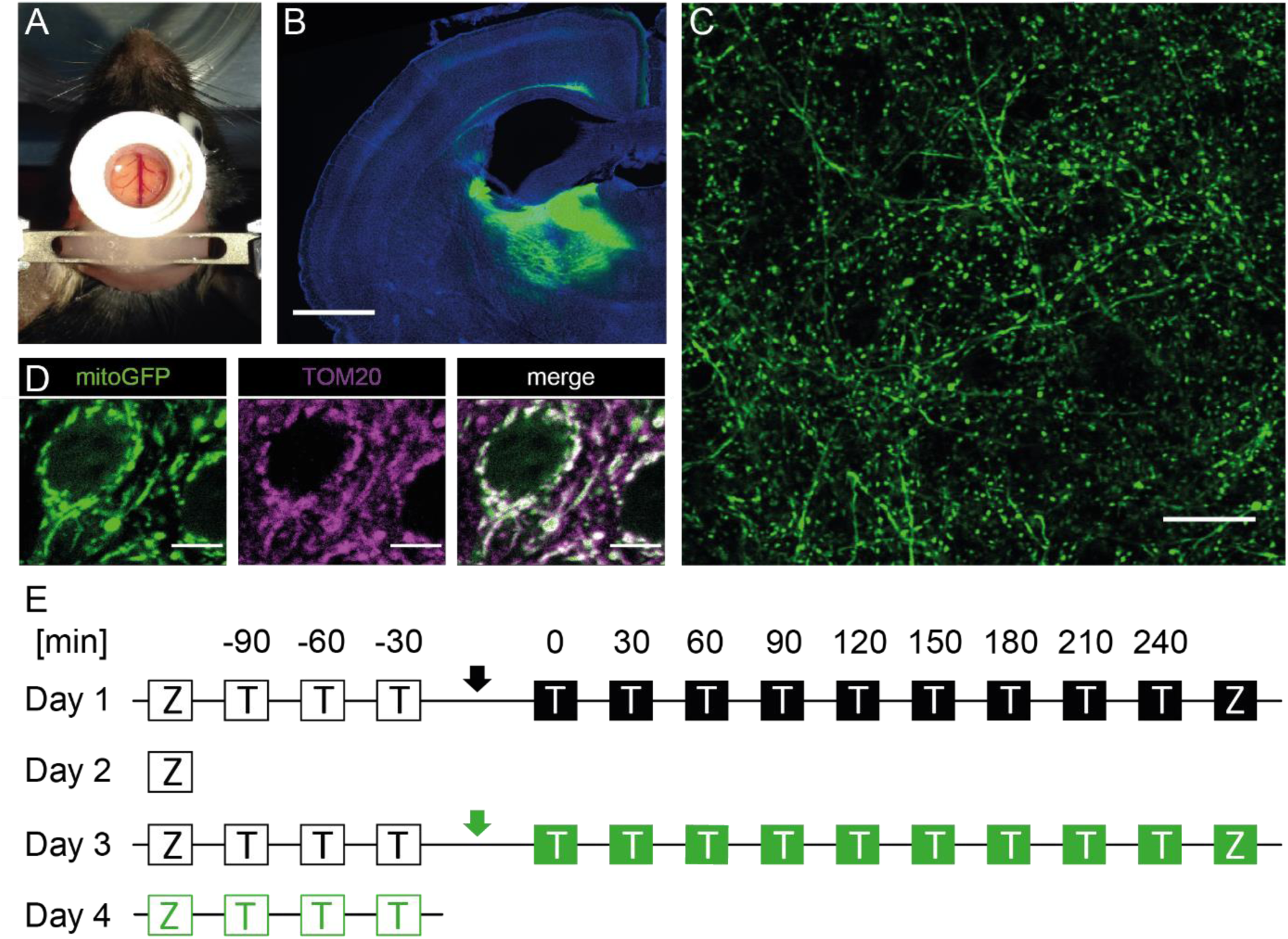
Visualizing axonal organelle trafficking under saline and ethanol conditions using *in vivo* two-photon microscopy. **(A)** Two-photon imaging was accomplished by implanting a chronic cranial window. The 3d-printed crown, needed for head-fixation of the animal, was attached to the glass coverslip. **(B)** Tile scan of thalamic injection site with projections reaching up to the cortex (wide field epifluorescence, scale bar 1 mm). **(C)** GFP-labelled mitochondria in thalamic projections imaged in upper layers of the cortex (two-photon, scale bar 25 µm). **(D)** Immunofluorescence staining of GFP-labelled mitochondria in neurons of the MD thalamus co-stained with the outer mitochondrial membrane marker TOM20 (single confocal plane, scale bar 5 µm). **(E)** For each mouse the experiment consisted of four imaging sessions. Baseline imaging (unfilled boxes) was acquired each day. On day 1 saline (black arrow) and on day 3 ethanol (green arrow) were administered i.p. followed by post-application imaging (filled boxes). On day 2 and 4, 24 h post-application images were recorded. Z = 3D-Stack, T = time lapse. Time lapses (10 min mitochondria; 5 min DCVs) were acquired every 30 min. 3D-stacks covered a total z distance of 50 µm with inter-plane distances of 1 µm.

Expression of fluorophore-tagged proteins labelling mitochondria (mitoGFP), DCVs (NPY-Venus) or presynaptic boutons (SyPhy-mCherry) was achieved by stereotaxically injecting rAAVs into the thalamus of mice (Figure 1B and C, suppl. Figure 1A, B for viral constructs). To verify the specificity of the marker proteins for tagging the desired organelles, we co-stained brain slices of injected mice with the outer mitochondrial membrane marker TOM20, the synaptic vesicle marker synaptophysin 1 or the endogenous DCV marker chromogranin-A (Chr-A, Knabbe et al. (2018)). For each marker, a co-localization of the fluorescently tagged proteins and the respective antibody was found (Figure 1D, suppl. Figure 1C). Time-lapse movies of mitochondria and DCVs as well as 3D-stacks before, 4 h and 24 h after i.p. injection of either saline or ethanol were acquired in anesthetized mice using two-photon microscopy (Figure 1E).

### Characteristics of mitochondrial transport *in vivo*

Figure 2A shows an exemplary axonal stretch representing the diverse facets of axonal mitochondrial transport such as transport speed, direction and temporary pausing. Kymographs were used to qualitatively and quantitatively describe mitochondrial transport events (Figure 2B).

**Figure 2:**
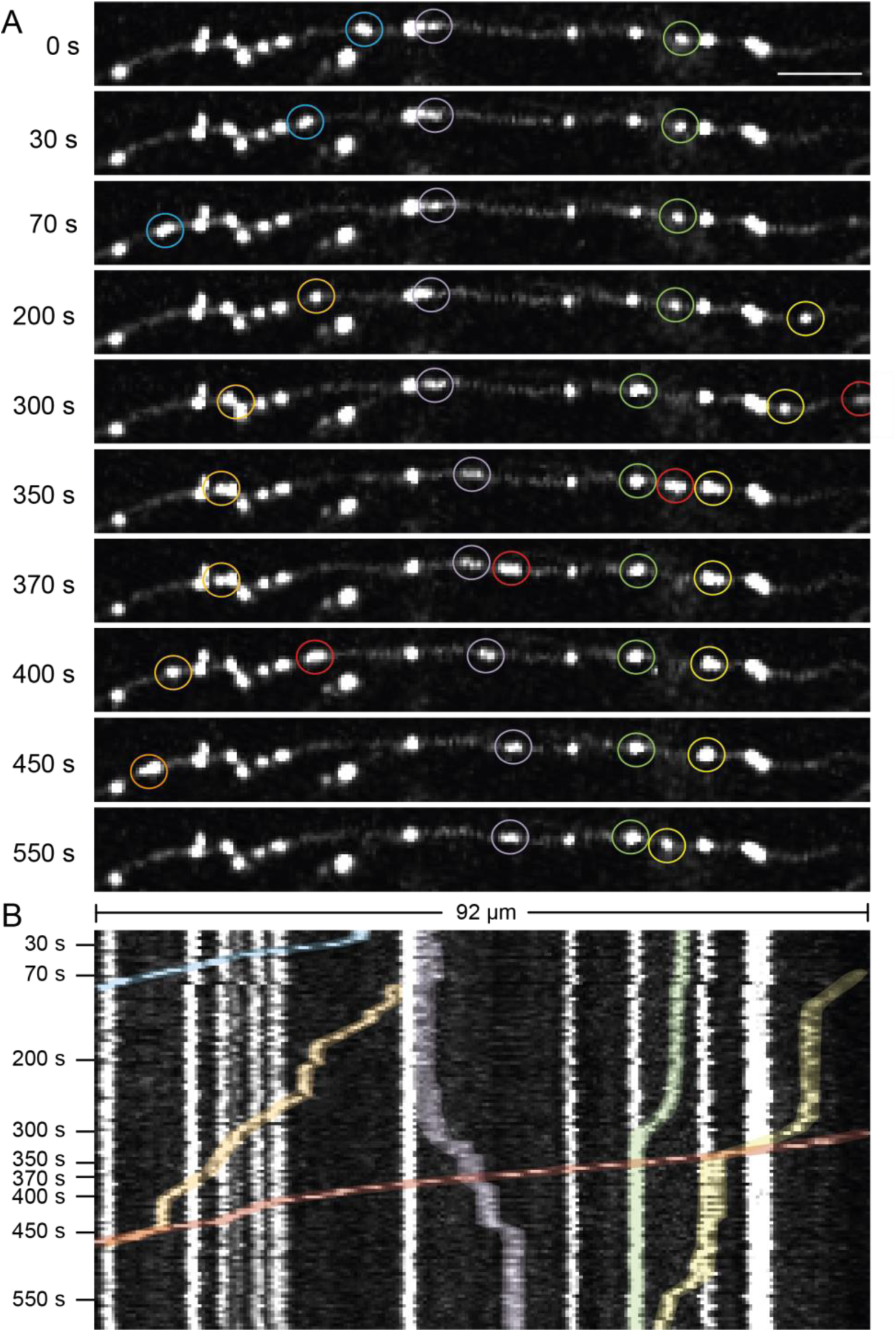
Qualitative description of characteristic features of axonal mitochondrial transport in the living mouse brain. **(A)** Axonal stretch at different time points during a time lapse of 600 s (two-photon, scale bar 10 μm). **(B)** Corresponding kymograph to axonal stretch in **A**. Colored mitochondria exemplify different features of mitochondrial transport. Orange: Slow mitochondrial trafficking. Red: Fast mitochondrial trafficking. Note that the red and orange mitochondria fused or started to co-migrate at approx. 450 s. Transport occurred in anterograde and retrograde directions as can be seen when comparing the purple mitochondrion to other colors. Blue: Remobilization of a stable mitochondrion at approx. 10 s. Purple: Relocation of a mitochondrion. Green: Fusion or co-localization of two mitochondria at approx. 300 s. Yellow: Slow mitochondrial transport with intermittent pauses (100 – 200 s) and temporary fusion/co-localization (350 – 550 s) with another mitochondrion. White: Immobile mitochondria.

We observed that mitochondrial transport was of variable speed: slow transport (orange) as well as fast transport (red) were detected. Further, mitochondrial trafficking occurred in anterograde as well as in retrograde direction, for example compare the purple mitochondrion with mitochondria of other colors. However, antero- and retrograde transport could not be distinguished, because the axons could not be traced back to the soma of the neurons due to their deep location in the thalamus. We could further observe that stationary or mobile properties were not fixed states, but mitochondria switched from one to the other. A stationary mitochondrion started to move at approximately 10 s (blue). We often found that mitochondria co-resided or co-migrated in the axon, during which they could possibly undergo fusion or fission (e.g. the green mitochondrion at 300 s). However, the resolution of two-photon microscopy did not allow distinguishing co-localization from fusion. Nonetheless, it can be assumed that fission/fusion occurred in co-residing or co-migrating mitochondria as mitochondrial fusion and fission events have been regularly described in axons (Cagalinec et al. 2013). The purple mitochondrion seemingly divided into smaller fractions which were relocated independently of each other, allowing to cover a greater length of the axon with a higher number of mitochondria or to equip more presynaptic boutons with individual mitochondria. Vice versa, smaller mitochondria, such as the green one, seemingly fused to a single large one. At times, mitochondria took pauses both at sites where other mitochondria resided or where no mitochondrion was yet located, e.g. the yellow mitochondrion. We could also detect a high number of stationary mitochondria that did not move during the time-lapse (white). In many cases, these could be found at the same location throughout subsequent time points, in rare cases even during the 4 imaging days (not shown).

First, we quantified mitochondrial transport under basal conditions. On average 8.47 ± 4.69 % of mitochondria were mobile (Figure 3A, n = 72 focal planes from 4 mice, Supp. Movie 1).

**Figure 3:**
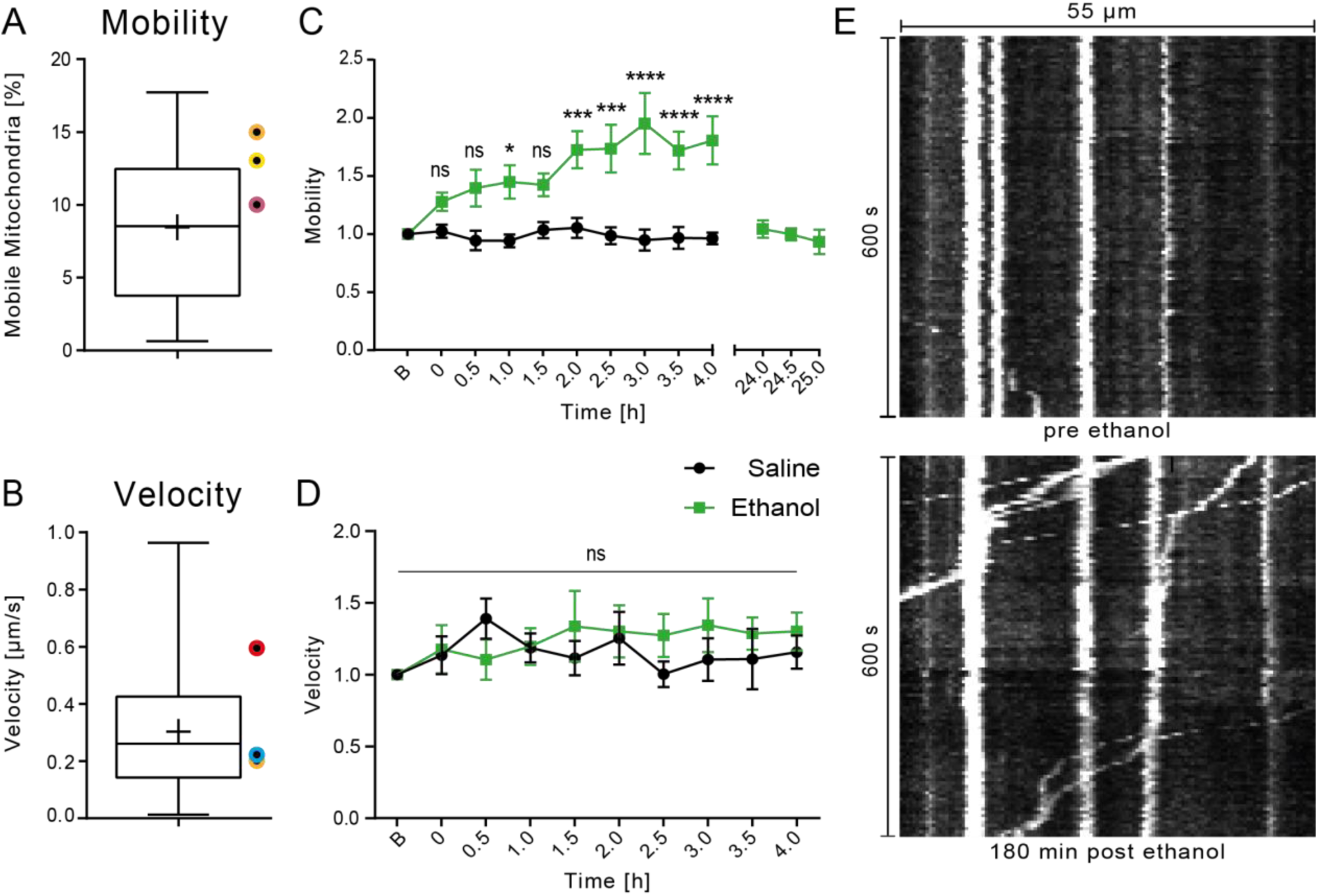
Mitochondrial mobility but not velocity increases after ethanol intoxication. **(A)** Under basal conditions 8.47 ± 4.69 % of mitochondria were mobile (n = 72 focal planes from 4 mice). Data displayed as Box-Whisker Plot; mean indicated as +, whiskers show min-max. Colored points indicate findings of previous studies. **(B)** Mean velocity under basal conditions was 0.30 ± 0.20 µm/s (n = 406 mitochondria from 4 mice). Data displayed as Box-Whisker Plot; mean indicated as +, whiskers show min-max. Colored points indicate findings of previous studies. **(C)** Time course of mitochondrial mobility under basal conditions (black) and after ethanol injection (green). Mitochondrial mobility increased up to two-fold during acute ethanol intoxication, but returned to baseline (B) after 24 h (n =11 focal planes from 4 mice; mean ± SEM). **(D)** Time course of mitochondrial velocity under basal conditions (black) and after ethanol injection (green). Mitochondrial velocity was unaffected by acute ethanol intoxication (n = 8 focal planes from 4 mice; mean ± SEM). **(E)** Kymograph before ethanol injection; only one mitochondrion was mobile. Kymograph of the same axonal stretch 180 min after ethanol injection. An increase in mitochondrial mobility can be seen. Two-way repeated measures ANOVA in **C, D** with post-hoc Sidak’s multiple comparisons test. ns = not significant, * = p<0.05, ** = p<0.005, *** = p<0.001, **** p<0.0001. Orange: Berthet et al. 2014; Yellow: Misgeld et al. 2007; Dark pink: Lewis et al. 2016; Red: Kiryu-Seo et al. 2010; Light blue: Takihara et al. 2015.

This is in line with previous reports as Lewis and co-authors (Lewis et al. 2016) reported 10 % moving mitochondria in axons of layer 2/3 cortical pyramidal neurons in awake as well as anesthetized mice. In addition, Misgeld et al. (2007) reported 13 % mobile mitochondria in axons of peripheral nerves in acute intercostal nerve-muscle explants of mice. *In vitro*, Berthet et al. (2014) found 15 % mobile mitochondria in axons of cultured hippocampal neurons of mice, though it has to be noted that in immature *in vitro* cultures mitochondria show higher motility than *in vivo* (Lewis et al. 2016). The recorded mean speed of axonal mitochondrial transport speed *in vivo* was 0.30 ± 0.20 µm/s (Figure 3B, n = 406 mitochondria of 4 mice). Other studies found average mitochondrial mean velocities ranging between approx. 0.20 µm/s (Takihara et al. (2015) *in vivo*; Berthet et al. (2014) *in vitro*) and 0.6 µm/s (Kiryu-Seo et al. 2010, *in vitro*). The smallest mean speed we observed was 0.01 µm/s, while the largest observed mean speed was 1.18 µm/s. To address the high standard deviation of the mean mitochondrial transport velocity, we calculated a frequency distribution of mitochondrial mean speeds (Supp. Fig. 2). Values were not normally distributed, however, a sum of two Gaussian distributed populations with mean1 = 0.16 ± 0.11 µm/s and mean2 = 0.40 ± 0.20 µm/s fitted the frequency distribution with R² = 0.9578, arguing for two distinct subsets of mitochondria that were transported at lower or faster speeds, respectively.

### Increased mitochondrial mobility after acute ethanol intoxication

To reveal the effects of acute ethanol intoxication on mitochondrial transport *in vivo*, we first imaged cortices every 30 min for 1.5 h to determine how mitochondria were transported under baseline conditions. The mice then received an i.p. injection of saline or ethanol, resulting in a blood alcohol concentration of approximately 250 mg/dl (Eisenhardt et al. 2015). After the injection, we continued to image the same brain regions every 30 min for the following four hours, as well as three times 24 h after ethanol. With regard to mitochondrial mobility, we found that over time mitochondrial mobility increased significantly in the ethanol conditions compared to the saline control condition (F (9, 90) = 3.517, p = 0.0009, n = 11 focal planes from 4 mice, two-way repeated measures ANOVA, Figure 3C, Supp. Movie 2). We observed that acute ethanol intoxication significantly increased mitochondrial mobility after 60 min (p = 0.0221; Sidak’s correction) peaking at a two-fold increase after 180 min (p < 0.0001; Sidak’s correction). Mitochondrial mobility remained highly increased even 180-240 min after the ethanol injection when ethanol is already expected to be eliminated from the blood (Eisenhardt et al. 2015). 24 h after intoxication, mitochondrial mobility had returned to baseline levels. In contrast to mitochondrial mobility, mitochondrial velocity remained unchanged after acute ethanol intoxication (F (9, 126) = 0.9890, p = 0.4525, n = 8 focal planes from 4 mice, two-way repeated measures ANOVA, Figure 3D). Representative kymographs (Figure 3E) of the same axonal stretch before and 180 min after ethanol application depict the increase in mitochondrial mobility during acute ethanol intoxication. Before ethanol administration, only one mitochondrion passed the axonal stretch and four mitochondria were stationary. Conversely, 180 min after ethanol intoxication the number of stationary mitochondria had not changed but seven mitochondria were transported. In conclusion, acute ethanol application increased mitochondrial mobility without affecting the transport velocity.

### DCV movement characteristics

To investigate whether the increase in mitochondrial transport mobility after acute ethanol intoxication was specific for mitochondria or if it affected microtubule-based transport in general, we analyzed the movement characteristics of DCVs, which are predominantly transported via microtubules (Zahn et al. 2004). Analyses of saline control and ethanol conditions was performed with an automated tracking algorithm allowing for the subsequent characterization of thousands of mobile particles per time-lapse video (Supp. Movie 3). We found that on average 69.4 ± 7.16 % of DCVs were mobile under basal conditions (n = 18 FOV from 3 mice, Figure 4A) which is in good agreement to a study by de Wit et al. (2006) who found 75.1 % moving DCVs in axons of cultured mouse cortical neurons.

**Figure 4:**
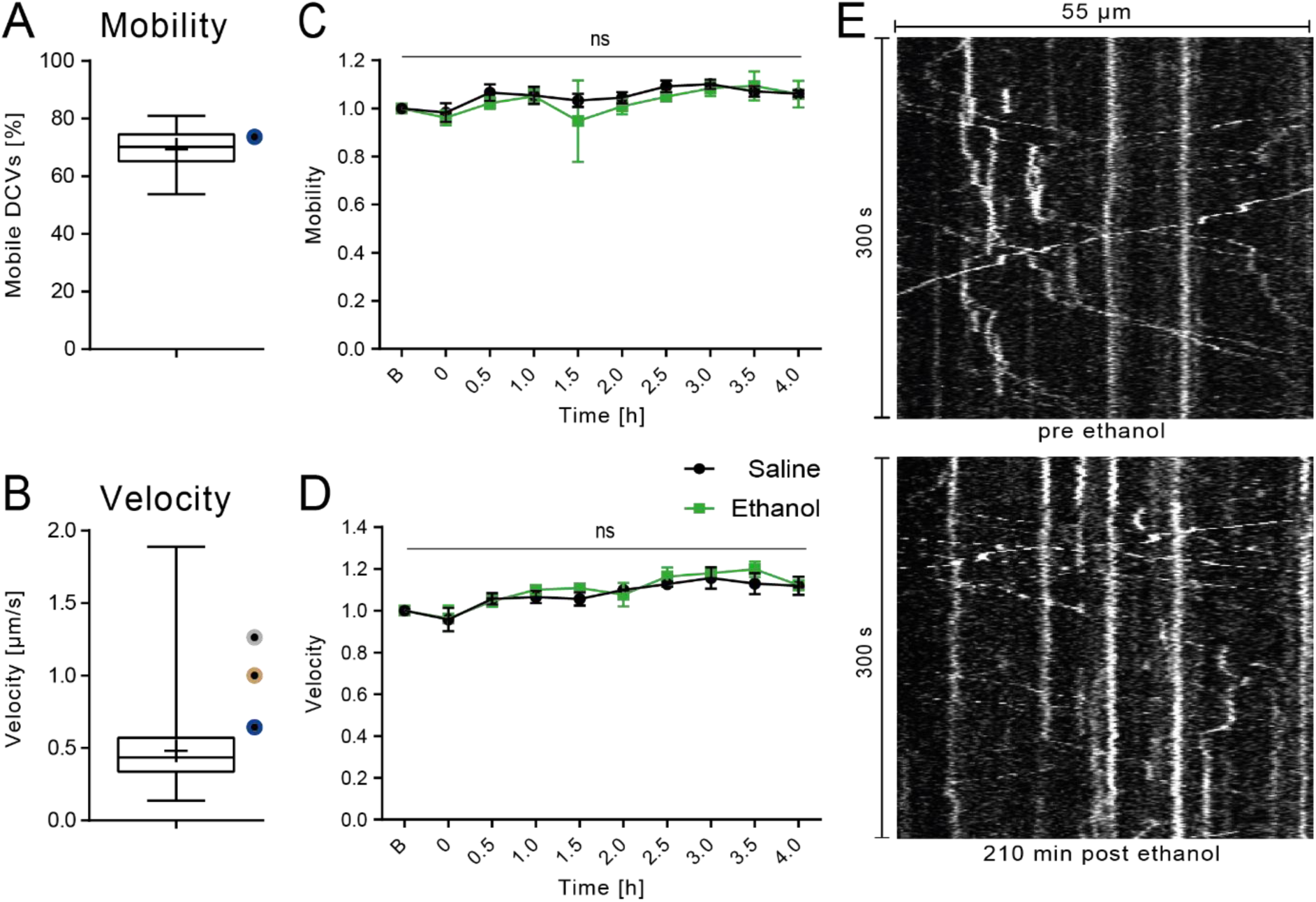
DCV transport remains stable after ethanol intoxication. **(A)** Under basal conditions 69.4 ± 7.16 % of DCVs were mobile (n = 18 FOV from 3 mice). Data displayed as Box-Whisker Plot; mean indicated as +, whiskers show min-max. Colored points indicate findings of previous studies. **(B)** Mean velocity under basal conditions was 0.48 ± 0.2 µm/s (n = 4657 DCVs from 3 mice). Data displayed as Box-Whisker Plot; mean indicated as +, whiskers show min-max. Colored points indicate findings of previous studies. **(C)** Time course of DCV mobility under basal conditions (black) and after ethanol injection (green). DCV mobility did slightly increase after acute ethanol intoxication as well as in the saline condition, the increase was not significantly different between saline and ethanol injected mice (n = 3 mice, mean ± SEM). **(D)** Time course of DCV velocity under basal conditions (black) and after ethanol injection (green). DCV velocity did increase after acute ethanol intoxication as well as in the saline condition, the increase was not significantly different between saline and ethanol injected mice (n = 3, mean ± SEM). **(E)** Kymographs of the same axonal stretch before and 210 min post-ethanol injection. The number of transported DCVs remains stable. Linear regression in **C, D.** ns = not significant. Dark blue: Wit et al. 2006; Grey: Kwinter et al. 2009; Brown: Knabbe et al. 2018.

The recorded mean speed of axonal DCV transport *in vivo* was 0.48 ± 0.2 µm/s (Figure 4B) (n = 4657 DCVs). A previous report from our laboratory (Knabbe et al. 2018) described a mean velocity of 1.03 µm/s in thalamic axons of anaesthetized mice. *In vitro*, values of 1.27 µm/s and 0.61 µm/s were reported in axons of cultured hippocampal and cortical neurons, respectively (Kwinter et al. 2009, de Wit et al. 2006).

### DCV mobility and velocities did not change after acute ethanol intoxication

In contrast to mitochondrial transport, DCV transport mobility was not affected by ethanol intoxication as could be shown by linear regression analyses where saline and ethanol slopes were not significantly different from each other (F (1, 16) = 0.247336, p = 0.6257 for slope equality, F(1, 17) = 2.56873, p = 0.1274 for intercepts equality). Figure 4C shows that in the saline control and ethanol condition, the mean DCV mobility increased only slightly over the time course of the experiment. DCV velocity increased by 20 % after 3.5 h (Figure 4D). This was seen in the saline condition as well, implicating an effect of the prolonged isoflurane anesthesia on DCV mobility. These results suggest that acute ethanol had no measurable effect on DCV transport velocity either (linear regression analysis, (F (1, 16) = 0.372645, p = 0.5501 for slope equality, F (1, 17) = 1.37549, p = 0.257 for intercepts equality). DCV transport is visualized in representative kymographs in Figure 4E where the same axonal stretch before and 210 min after ethanol application is shown. Both kymographs reveal comparable numbers of transported and stationary DCVs.

### Effects of higher mitochondrial mobility on mitochondrial length and abundance

We next wanted to investigate whether the higher amount of mobile mitochondria after ethanol administration was caused (i) by re-mobilizing previously stationary mitochondria, (ii) by recruiting mitochondria from distant regions, e.g. the soma, or (iii) by stable mitochondria undergoing fission. Therefore, mitochondrial length and axonal abundance before, 4 h and 24 h after ethanol injection (Figure 5A) was characterized.

**Figure 5:**
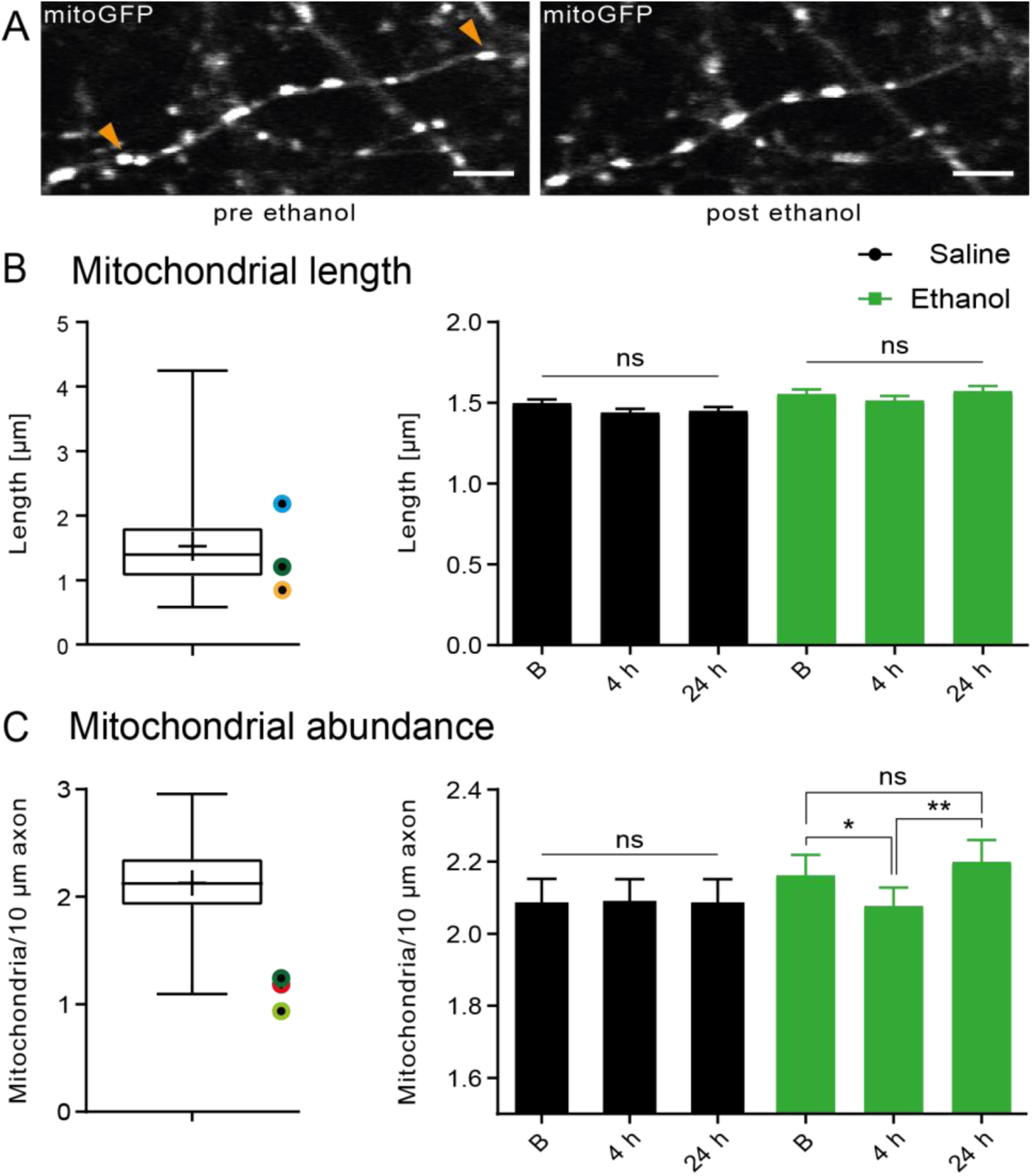
Acute ethanol intoxication does not affect mitochondrial structure, but acutely decreases mitochondrial abundance. **(A)** Representative images of delocalized mitochondria (orange arrow heads) 4 h after ethanol application (two-photon, scale bar 5 µm). **(B)** Mean mitochondria length was 1.52 ± 0.73 µm (n = 1343 mitochondria). Data displayed as Box-Whisker Plot; mean indicated as +, whiskers show 1-99 percentile. Colored points indicate findings of previous studies. Mitochondrial length (n > 600 mitochondria per time point from 4 mice, mean ± SEM) was unaffected by acute ethanol intoxication. **(C)** On average 2.128 ± 0.3648 mitochondria were found per 10 µm axon (n = 80 axons from 4 mice). Data displayed as Box-Whisker Plot; mean indicated as +, whiskers show min-max. Colored points indicate findings of previous studies. 4 h after ethanol intoxication mitochondrial abundance was significantly reduced (n = 80 axons from 4 mice), but returned to baseline (B) levels again after 24 h. Kruskal-Wallis Test with post-hoc Dunn‘s multiple comparisons test in **B.** Two-way repeated measures ANOVA with Sidak’s multiple comparisons test and **C**. ns = not significant, * = p<0.05, ** = p<0.01. Orange: Berthet et al. 2014; Dark green: Chang et al. 2006; Red: Kiryu-Seo et al. 2010; Light blue: Takihara et al. 2015; Light green: Smit-Rigter et al. 2016.

The mean mitochondrial length was 1.52 ± 0.73 µm (n = 1343 mitochondria, Figure 5 B left). Other studies reported values of 0.77 µm in axons of dopaminergic neurons in the nucleus accumbens and caudate putamen of mice (Berthet et al. 2014), 1.4 µm in axons of cultured cortical neurons of rats (Chang et al. 2006) and 2.6 µm in moving mitochondria of retinal ganglion cells in mice (Berthet et al. 2014, Takihara et al. 2015). Ethanol injection did not change mitochondrial length suggesting that an increased number of fission events was not responsible for more moving mitochondria (p > 0.9999 for pre saline vs. 4 h post saline, pre saline vs. pre ethanol, pre ethanol vs. 4 h post ethanol, pre ethanol vs. 24 h post ethanol and p = 0.5532 for pre saline vs. 24 h post saline, Kruskal-Wallis test with Dunn’s correction; Figure 5B right). Mitochondrial abundance was 2.128 ± 0.3648 mitochondria/10µm axon (n = 80 axons from 4 mice, Figure 5C left) resulting in an average inter-mitochondrial distance of 4.70 µm. Other studies reported values of 0.9 mitochondria/10 µm in axons of layer 2/3 pyramidal neurons in the primary visual cortex of mice (Smit-Rigter et al. 2016), 1.3 mitochondria/10 µm in axons of cultured cortical neurons of rats (Chang et al. 2006) and 1.2 mitochondria/10 µm in axons of cultured dorsal root ganglion cells of rats (Kiryu-Seo et al. 2010). Similarly to mitochondrial length, mitochondrial abundance can be expected to vary between brain regions and cell types, as mitochondrial morphology and distribution most likely adjust to the individual energy demands of cells (McCarron et al. 2013). Measuring mitochondrial abundance 4 h and 24 h post saline and ethanol, we found that mitochondrial abundance was significantly altered in the ethanol condition (F (2, 156) = 3.728, p = 0.0262, n = 40 axons from 4 mice, two-way repeated measures ANOVA, Figure 5B left). Specifically, we found that mitochondrial abundance was significantly reduced 4 h after ethanol injection (p = 0.0372, Sidak’s correction, Figure 5C right). After 24 h, mitochondrial abundance increased again compared to 4 h after ethanol (p = 0.0012, Sidak’s correction), but was not significantly different to pre ethanol levels (p = 0.6152, Sidak’s correction) suggesting that mobile mitochondria were recruited from the pool of previously stationary mitochondria during the acute phase of ethanol intoxication, but returned to pre ethanol levels after 24 h. However, it cannot be ruled out that additional mitochondria were recruited from the soma.

### Effects of higher mitochondrial mobility on occupancy of presynaptic boutons

As most stationary mitochondria are located at presynaptic boutons and are important during high synaptic energy demands (Verstreken et al. 2005), we sought to investigate mitochondrial occupancy of synapses. Presynaptic boutons were tagged using the presynaptic vesicle protein synaptophysin fused to mCherry (Figure 6A).

**Figure 6:**
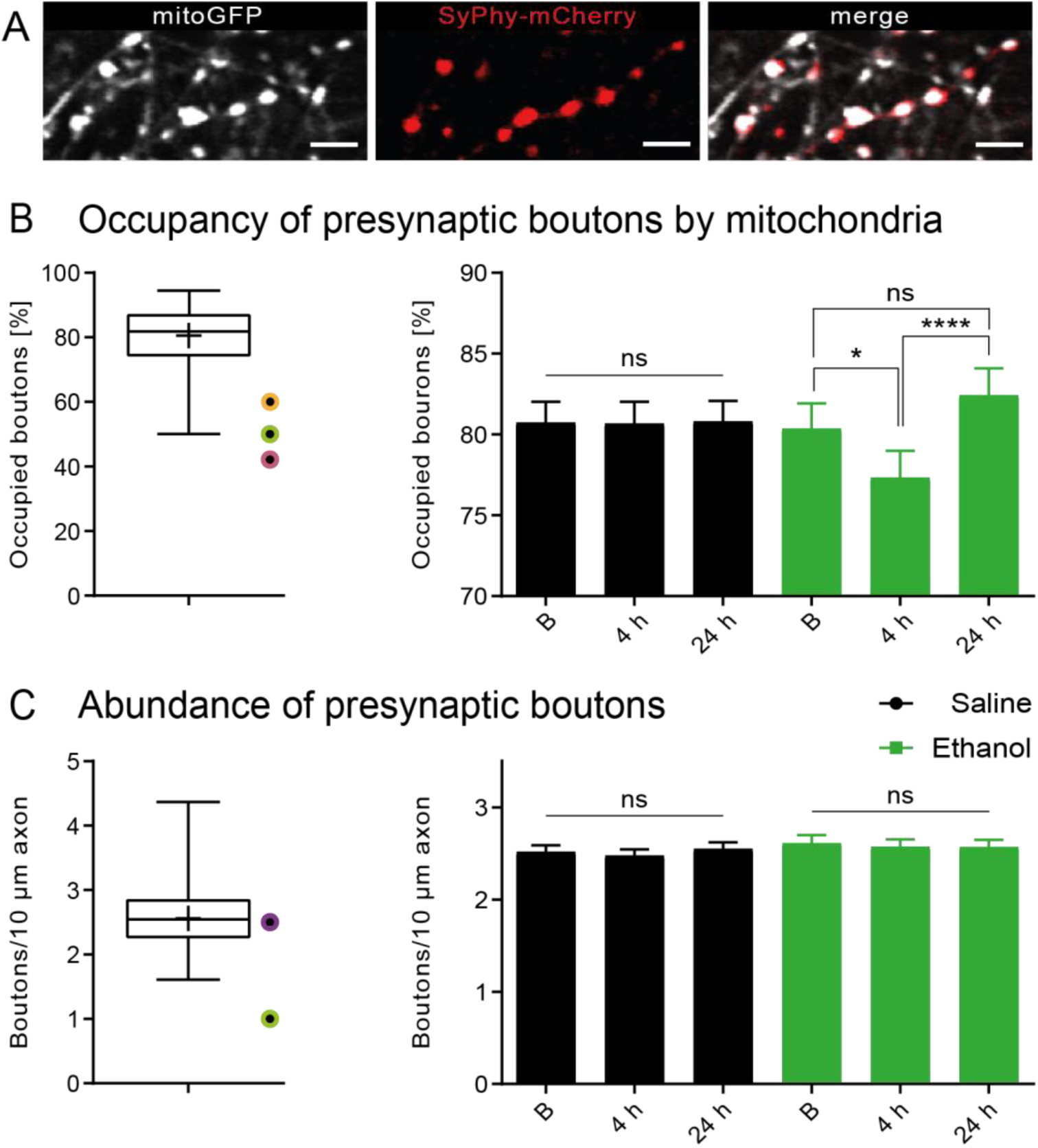
Acute ethanol intoxication decreases occupancy of presynaptic boutons by mitochondria, but does not affect presynaptic abundance. **(A)** Representative images of occupied as well as unoccupied presynaptic boutons. Mitochondria depicted in grey, boutons depicted in red (two photon, scale bar 5 µm). **(B)** On average 80.5 ± 9.1 % of presynaptic boutons (n = 80 axons from 4 mice) were occupied by mitochondria in thalamic long-projection neurons. Data displayed as Box-Whisker Plot; mean indicated as +, whiskers show min-max. Colored points indicate findings of previous studies. 4 h after ethanol intoxication occupancy of presynaptic boutons was significantly reduced. After 24 h occupancy returned to baseline level. **(C)** We found on average of 2.56 presynaptic boutons/10 µm axon (n = 80 axons from 4 mice) corresponding to an inter-synaptic distance of 4 µm. Data displayed as Box-Whisker Plot; mean indicated as +, whiskers show min-max. Colored points indicate findings of previous studies. Presynaptic abundance was unaffected by acute ethanol intoxication. Two-way ANOVA with Sidak’s multiple comparisons test in **B** and **C** (n = 40 axons from 4 mice). ns = not significant, * = p < 0.05, **** = p < 0.0001. Orange: Berthet et al. 2014; Light green: Smit-Rigter et al. 2016; Dark pink: Lewis et al. 2016; Purple: Becker et al. 2008.

The typically high copy number of synaptophysin on synaptic vesicles lead to a strong local fluorescence signal that allowed visualization of even small en-passant boutons. In contrast, other studies mostly characterized larger boutons that appeared as ellipsoid structures in axons labeled with cytosolic or membrane-bound fluorophores. We found that 80.5 ± 9.1 % of the presynaptic boutons on thalamic axons were occupied by mitochondria (n = 80 axons from 4 mice, Figure 6B left). Other studies report occupancy numbers of 42 % in axons of cultured cortical neurons (Lewis et al. 2016), 49 % in axons of layer 2/3 pyramidal neurons in the primary visual cortex of mice (Smit-Rigter et al. 2016) and approximately 60 % in axons of dopaminergic neurons in the nucleus accumbens and caudate putamen of mice (Berthet et al. 2014). In line with the decrease in mitochondrial abundance, we could show a decrease in mitochondrial occupancy of boutons in the ethanol condition (F (2, 156) = 5.382, p = 0.003, n = 40 axons from 4 mice, two-way repeated measured ANOVA, Figure 6B right). Presynaptic occupancy significantly decreased 4 h after ethanol administration (p = 0.0163, Sidak’s correction) and increased again after 24 h (p < 0.0001, Sidak’s correction) to a level that was similar to the pre ethanol level (p = 0.1574, Sidak’s correction) supporting the notion that during the acute phase of ethanol intoxication, mitochondria were recruited from a stationary pool of presynaptically localized mitochondria. The abundance of presynaptic boutons was 2.56 ± 0.52/10 µm resulting in an inter-synaptic distance of 3.9 ± 0.66 µm. Synaptic abundance seems to be largely dependent on the brain area of interest as Berthet et al. (2014) found 6 synapses/10 µm in axons of dopaminergic neurons in the nucleus accumbens and caudate putamen of mice, while Smit-Rigter et al. (2016) reported 1 synapse/10 µm in axons of layer 2/3 pyramidal neurons in the primary visual cortex of mice, and Becker et al. (2008) found 2.5 synapses/10 µm in hippocampal CA3 axons in mouse organotypic slice cultures. We now report that the inter-synaptic distance was not significantly altered after ethanol injection (F (2, 156) = 1.656, p = 0.1943, n = 40 axons from 4 mice, two-ways repeated measures ANOVA, Figure 6C right).

### Effects of ethanol injection on turnover of presynaptic boutons

As we did not see significant ethanol-mediated effects on the abundance of presynaptic boutons, we defined smaller, more specific subgroups of mitochondrially occupied and unoccupied presynaptic boutons. In these two subgroups, newly formed or lost presynaptic boutons were compared between two time points (pre injection vs. 4 h, pre injection vs. 24 h, Figure 7A).

**Figure 7:**
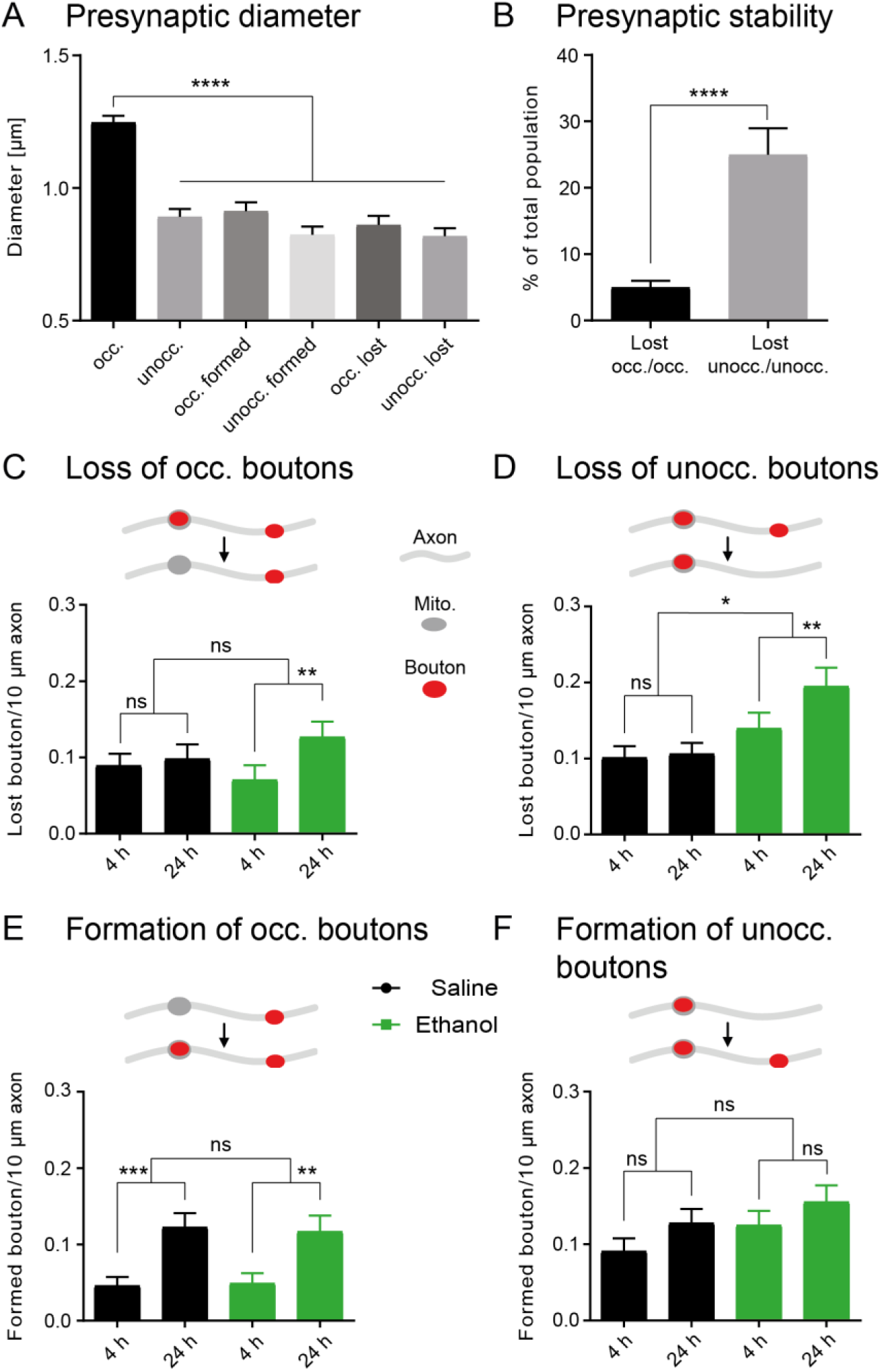
Occupied presynaptic boutons are larger and highly stable, while unoccupied presynaptic boutons are smaller and more susceptible for ethanol-induce demise. **(A)** Diameter of stable, lost and formed occupied (occ.) and unoccupied (unocc.) presynaptic boutons (n > 50 per group). Occupied stable presynaptic boutons were significantly larger than other boutons. Occupied boutons had on average a larger diameter (1.26 µm) than unoccupied boutons (0.89 µm) and also reached higher maximal diameters (3.03 µm) compared to unoccupied synapses (2.372 µm). **(B)** Stability of presynaptic boutons differed significantly in dependence of occupancy. **(C)** Loss of occupied presynaptic boutons was unaffected by acute ethanol intoxication when comparing the saline to the ethanol condition, however, a significant increase after 24 h compared to 4 h could be detected in the ethanol condition, but not in the saline condition. **(D)** Loss of unoccupied presynaptic boutons was significantly increased after 24 h by acute ethanol intoxication indicating that boutons devoid of mitochondria were more susceptible for ethanol-induce demise. **(E)** Formation of occupied presynaptic boutons was unaffected by ethanol intoxication. **(F)** Formation of unoccupied presynaptic boutons was unaffected by ethanol intoxication. Mann-Whitney tests in **A** and **B**. Two-way repeated measures ANOVA with post-hoc Sidaks‘s multiple comparisons test in **C-F** (n = 40 axons from 4 mice). ns = not significant, * = p < 0.05; ** = p < 0.01, *** = p < 0.001, **** = p < 0.0001

Analyzing the diameter of presynaptic boutons, we found that occupied stable boutons (1.25 ± 0.48 µm) were roughly 50% larger than unoccupied (0.89 ± 0.3 µm) or newly formed occupied boutons (0.91 ± 0.24 µm; p < 0.0001 for stable occupied boutons compared with all other groups, Mann-Whitney test, n > 50 per group). To elucidate whether stability of presynaptic boutons was dependent on occupancy by mitochondria, we calculated the fraction of lost boutons of either occupied or unoccupied boutons (Figure 7B). We found a significantly higher loss of unoccupied compared to occupied boutons (24.9 ± 25.5 % vs. 5 ± 6.3 %, p < 0.0001, Mann Whitney test), confirming the finding of Smit-Rigter et al. (2016) that mitochondria play a pivotal role for the stability of presynaptic boutons *in vivo*. Finally, we wanted to investigate if ethanol affects the loss or formation of mitochondrially occupied or unoccupied boutons. In occupied boutons, we found no significant effect of ethanol on the loss rate (F (1, 78) = 0.05087, p = 0.8221, n = 40 axons from 4 mice, two-way repeated measures ANOVA, Figure 7C), though there was a trend towards an increase in the loss of occupied boutons between 4 h and 24 h after ethanol exposure that was not seen in the saline condition. In unoccupied boutons, ethanol significantly increased the rate of loss compared with the saline condition (F (1, 78) = 6.512, p = 0.0127, n = 40 axons from 4 mice, Figure 7D). This effect was significantly higher 24 h after ethanol compared to 4 h (p = 0.007, Sidak’s correction). We did not see a significant effect of ethanol on the formation of occupied (F (1, 78) = 0.004360, p = 0.9475, n = 40 axons from 4 mice, two-way repeated measures ANOVA, Figure 7E) or unoccupied boutons (F (1, 78) = 2.186, p = 0.1433, n = 40 axons from 4 mice, two-way repeated measures ANOVA, Figure 7F) compared to the saline condition. Formation of occupied boutons was significantly increased in both conditions after 24 h compared to 4 h (p = 0.0005 for saline, p = 0.0024 for ethanol, Sidak’s correction), suggesting that formation of presynaptic boutons starts with a presynaptic vesicle cluster and mitochondria follow for stabilization. In conclusion, we found that occupied presynaptic boutons are larger than unoccupied boutons and that unoccupied boutons are more likely to be lost under the acute influence of ethanol.

## Discussion

*In vivo*, longitudinal two-photon time-lapse imaging was used to visualize genetically labeled presynaptic boutons, mitochondria and DCVs in mildly anesthetized mice. In the living brain, acute ethanol intoxication changed the turnover of presynaptic boutons devoid of mitochondria and increased the mobility of mitochondria, while DCV trafficking remained unaffected. Based on the rapid onset of ethanol action on mitochondrial mobility (Figure 3C) we propose that presynaptic boutons have an increased probability of losing mitochondria during a 4 h period after ethanol administration (Figure 6B), in turn rendering them more susceptible to turnover and loss (Figure 7D). This novel mode of ethanol action may explain memory deficits frequently observed in ethanol intoxication (Zorumski, Mennerick and Izumi 2014).

Under control conditions, mitochondrial movement dynamics in our study were similar to other *in vivo* studies (Misgeld et al. (2007) Lewis et al. (2016)), confirming their finding of reduced mitochondrial movement dynamics when compared to *in vitro* observations (Lewis et al. 2016). We now find that ethanol almost doubled the initial mobility after 3 h, with the effect lasting longer than four hours. We can only speculate why mitochondrial mobility was increased after ethanol intoxication in mice. In mice, metabolic ethanol elimination is completed within about three hours (Eisenhardt et al. 2015). This argues against the possibility that ethanol directly affects adaptor or motor proteins, as the increase in mitochondrial mobility remained high after four hours when ethanol should have been eliminated. DCV transport was unaffected by acute ethanol intoxication, although many transport and adaptor proteins of DCV transport are similar to those in mitochondrial transport, rendering a direct effect of ethanol on these proteins unlikely. One possible explanation might be that ethanol increases GSK3Beta-dephosphorylation at Serin 9 (Liu et al. 2009), thereby increasing the activity of GSK3Beta. Overexpression of GSK3Beta was shown to increase trafficking of mitochondria in neurons (Llorens-Martin et al. 2011).

Analysis of DCV trafficking was performed by a fully automated tracking algorithm, allowing us to analyze roughly 180000 individual tracks over time from about 45 gigabyte of data. The mean DCV speed reported here (0.48 ± 0.2 µm/s) was lower compared to our previous work (1.03 ± 0.03 µm/s in Knabbe et al. (2018)) where we specifically focused on investigating axons exhibiting a large number of trafficking events. As the imaging speed was relatively slow, faster vesicles were difficult to track for the algorithm, possibly creating a bias towards slower trafficking speed. Nonetheless, this bias does not constitute a problem, because all DCV trafficking was analyzed with the same approach. The analysis revealed a possible effect of the isoflurane anesthesia on trafficking, though no specific effect of ethanol on DCV mobility, contrasting with previously published work (Iacobucci and Gunawardena 2018). This discrepancy might be due to differences in model systems. *Drosophila melanogaster* larvae were immersed in alcoholic solution in the other study, potentially allowing higher amounts of alcohol to reach the neurons. Furthermore, as Iacobucci and Gunawardena (2018) described, *Drosophila* larvae naturally live in and feed on fermented fruit and therefore have to cope with alcohol in their direct environment, possibly explaining differences in alcohol sensitivity of organelle trafficking. In conclusion, we interpret the differential effect of ethanol on mitochondrial and DCV transport to indicate that ethanol does not directly act on the microtubular transport machinery for organelles.

We consider the relationship of mitochondrial mobility and occupancy of presynaptic boutons in the context of ethanol action a promising new mechanism that may even help to understand memory loss observed after acute alcohol intoxication. In contrast to previous studies mostly focusing on large axonal ellipsoid structures (Smit-Rigter et al. 2016, Becker et al. 2008), we also included the analysis of small synaptic vesicle clusters, thereby covering the entire range of presynaptic boutons sizes. We found that the mitochondrial occupancy of presynaptic boutons strongly decreased within 4 h, yet mitochondrial length, an indicator of the dynamics of mitochondrial fusion and fission, was unaltered after ethanol injection. This may suggest that previously stable mitochondria were mobilized from presynaptic boutons rather than mitochondrial fission creating smaller units. The mitochondrial occupancy of around 80% of all presynaptic boutons was relatively high compared to other studies, which might be specific for the long range thalamic projections that we analyzed, where synapses might form as far as 3 mm away from the cell soma. On the basis of the high mitochondrial occupancy, one could speculate that long-range projections tend to produce more stable synapses. Alternatively, a technical explanation is that we covered a substantial range of bouton sizes and therefore also sampled smaller structures that were not considered in previous *in vivo* imaging studies.

Upon characterization of presynaptic boutons in the saline condition, we could show that occupied stable boutons were significantly larger than newly formed or unoccupied boutons. The number of occupied boutons significantly increased in the saline and the ethanol conditions after 24 h compared to 4 h. This may suggest that newly formed presynaptic boutons initially start as synaptic vesicle clusters that grow by recruiting mitochondria, thereby stabilizing the nascent bouton by meeting its energy demands (Smith et al. 2016). When we analyzed the fraction of lost synapses that were occupied before and compared those to lost unoccupied synapses, we found that the latter were highly unstable. Every fourth presynaptic unoccupied bouton was lost four hours after the first imaging. This potentially underlines the importance of stabilization by mitochondria of newly formed boutons. When this step does not occur, presynaptic boutons might have a higher probability to vanish. This is in agreement with a study from Qiao et al. (2016) who could show that newly formed synaptic boutons perished significantly more often in the next 2-3 weeks than pre-existing synaptic boutons. Thus, it could be speculated that mitochondria have an influence on whether a bouton vanishes again after a few days to weeks or becomes part of a synapse that may persist for months.

Ethanol intoxication led to a loss of unoccupied boutons, perhaps because those presynaptic boutons were not yet in the process of becoming stable presynaptic boutons and were as such more susceptible to an ethanol-induced demise. Processes such as learning and memory in the adult brain require plastic changes including formation of new synapses (Knott et al. 2002) or elimination and remodeling (weakening or strengthening) of already existing synapses (Lamprecht and LeDoux 2004). This effect of ethanol on mostly unoccupied presynaptic boutons could explain why memory formation becomes increasingly difficult with higher amounts of ethanol and why newly formed memories might not be as stable after or during an excessive consumption of ethanol.

Thus, newly formed presynaptic boutons were less likely to be stabilized after ethanol intoxication, maybe due to the increased amount of trafficked mitochondria which might be recruited from synapses that were just in the process of being stabilized.

In summary, we could identify mitochondrial transport as a novel target of acute ethanol intoxication. The increase in mitochondrial mobility led to a redistribution of mitochondria along the axon resulting in a reduction of presynaptic occupancy by mitochondria after 4 h. As a result of the ethanol intoxication, more unoccupied presynaptic boutons perished after 24 h. We were able to show that occupied presynaptic boutons were larger in size as well as more stable than unoccupied presynaptic boutons, but might also be slightly affected by ethanol intoxication. We therefore suggest that mitochondria have a positive effect on the stability of presynaptic boutons and that the redistribution of mitochondria during acute ethanol intoxication preferentially renders unoccupied presynaptic boutons more susceptible to ethanol-induced demise. By revealing *in vivo* effects of acute ethanol on mitochondrial trafficking and stability of presynaptic boutons, we provide a possible explanation for the well-known observation that learning and memory are impaired after consuming high amounts of ethanol.

## Acknowledgments

We want to thankfully acknowledge the technicians Michaela Kaiser and Claudia Kocksch for the preparation of rAAVs. The data storage service SDS@hd supported by the Ministry of Science, Research and the Arts Baden-Württemberg (MWK) and the German Research Foundation (DFG) through the grant INST 35/1314-1 FUGG is gratefully acknowledged. We also acknowledge the support of the DFG within the SFB 1129.

## Contributions

J.P. performed all experiments, analyzed data and wrote the manuscript. A.J. and K.R. developed the automated tracking algorithm and analyzed data. S.C. gave the inspiration for the project and kindly provided viral constructs. J.K., J.P. and T.K. wrote the manuscript. J.K. devised the project, analyzed data and wrote the manuscript. All authors reviewed and approved the manuscript.

---

**Suppl. Figure 1:**
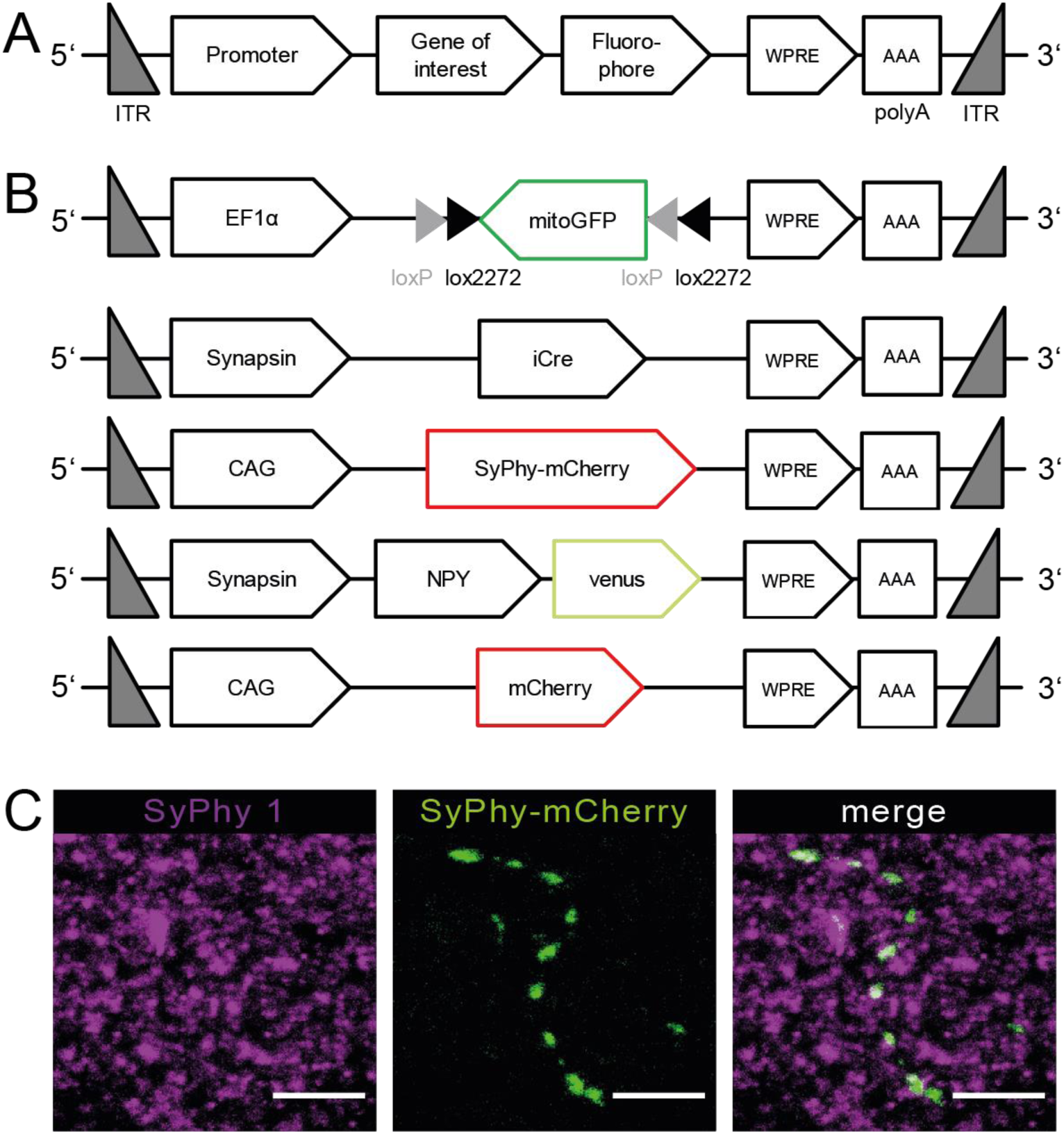
Viral constructs and verification of viral specificity. **(A)** Schematic structure of AAV constructs. The insert was flanked with inverted terminal repeats (ITR). Gene of interest and fluorophore expression was driven with a promoter. At the 3’ end the insert contained the woodchuck hepatitis post-translational element (WPRE) sequence and the bovine growth hormone poly-A sequence (polyA). **(B)** The constructs of EF1α-DIO-mitoGFP, Synapsin-iCre, CAG-SyPhy-mCherry, Synpasin-NPY-venus, Synapsin-BPY-mCherrry and CAG-mCherry. **(C)** Example image of SyPhy-mCherry co-stained with SyPhy 1 (MIP of six consecutive confocal planes, scale bar 5 µm). Verification of NPY-venus co-stained with Chromogranin-A can be found in Knabbe et. al., 2018.

**Suppl. Figure 2:**
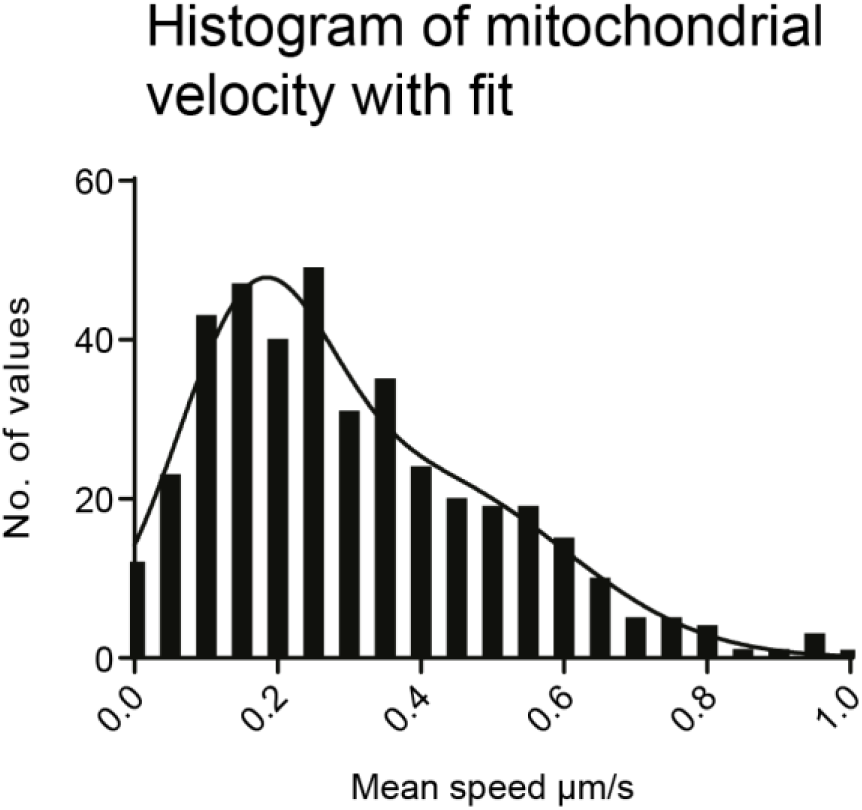
Frequency distribution of mitochondrial mean speeds. It could be shown that values were not normally distributed, however, a sum of two Gaussian distributed populations with mean1 = 0.16 ± 0.11 µm/s and mean2 = 0.40 ± 0.20 µm/s fitted the frequency distribution with R² = 0.9578 (n = 406 mitochondria from 4 mice), arguing for two distinct subsets of mitochondria which are transported at lower or faster speeds.

**Suppl. Movie 1: Mitochondrial trafficking in vivo under baseline conditions. Imaging framerate 0.2 Hz, sped up to 12 frames/s**.

**Suppl. Movie 2: Mitochondrial trafficking in vivo after ethanol injection. Imaging framerate 0.2 Hz, sped up to 12 frames/s**.

**Suppl. Movie 3: Dense core vesicle trafficking in vivo with automated tracking overlay. Imaging framerate 0.94 Hz, sped up to 25 frames/s**.

## Literature

Abrahao, K. P., A. G. Salinas & D. M. Lovinger (2017) Alcohol and the Brain: Neuronal Molecular Targets, Synapses, and Circuits. Neuron, 96, 1223–1238.

Becker, N., C. J. Wierenga, R. Fonseca, T. Bonhoeffer & U. V. Nagerl (2008) LTD induction causes morphological changes of presynaptic boutons and reduces their contacts with spines. Neuron, 60, 590–7.

Berthet, A., E. B. Margolis, J. Zhang, I. Hsieh, J. Zhang, T. S. Hnasko, J. Ahmad, R. H. Edwards, H. Sesaki, E. J. Huang & K. Nakamura (2014) Loss of mitochondrial fission depletes axonal mitochondria in midbrain dopamine neurons. J Neurosci, 34, 14304–17.

Bharat, V., M. Siebrecht, K. Burk, S. Ahmed, C. Reissner, M. Kohansal-Nodehi, V. Steubler, M. Zweckstetter, J. T. Ting & C. Dean (2017) Capture of Dense Core Vesicles at Synapses by JNK-Dependent Phosphorylation of Synaptotagmin-4. Cell Rep, 21, 2118–2133.

Billups, B. & I. D. Forsythe (2002) Presynaptic mitochondrial calcium sequestration influences transmission at mammalian central synapses. J Neurosci, 22, 5840–7.

Bodhinathan, K. & P. A. Slesinger (2013) Molecular mechanism underlying ethanol activation of G-protein-gated inwardly rectifying potassium channels. Proc Natl Acad Sci U S A, 110, 18309–14.

Boffi, J. C., J. Knabbe, M. Kaiser & T. Kuner (2018) KCC2-dependent Steady-state Intracellular Chloride Concentration and pH in Cortical Layer 2/3 Neurons of Anesthetized and Awake Mice. Front Cell Neurosci, 12, 7.

Cagalinec, M., D. Safiulina, M. Liiv, J. Liiv, V. Choubey, P. Wareski, V. Veksler & A. Kaasik (2013) Principles of the mitochondrial fusion and fission cycle in neurons. J Cell Sci, 126, 2187–97.

Cardoso, R. A., S. J. Brozowski, L. E. Chavez-Noriega, M. Harpold, C. F. Valenzuela & R. A. Harris (1999) Effects of ethanol on recombinant human neuronal nicotinic acetylcholine receptors expressed in Xenopus oocytes. J Pharmacol Exp Ther, 289, 774–80.

Chang, D. T., A. S. Honick & I. J. Reynolds (2006) Mitochondrial trafficking to synapses in cultured primary cortical neurons. J Neurosci, 26, 7035–45.

Chen, Y. & Z. H. Sheng (2013) Kinesin-1-syntaphilin coupling mediates activity-dependent regulation of axonal mitochondrial transport. J Cell Biol, 202, 351–64.

de Wit, J., R. F. Toonen, J. Verhaagen & M. Verhage (2006) Vesicular trafficking of semaphorin 3A is activity-dependent and differs between axons and dendrites. Traffic, 7, 1060–77.

Denk, W., K. R. Delaney, A. Gelperin, D. Kleinfeld, B. W. Strowbridge, D. W. Tank & R. Yuste (1994) Anatomical and functional imaging of neurons using 2-photon laser scanning microscopy. J Neurosci Methods, 54, 151–62.

Dopico, A. M., A. N. Bukiya & G. E. Martin (2014) Ethanol modulation of mammalian BK channels in excitable tissues: molecular targets and their possible contribution to alcohol-induced altered behavior. Front Physiol, 5, 466.

Dopico, A. M. & D. M. Lovinger (2009) Acute alcohol action and desensitization of ligand-gated ion channels. Pharmacol Rev, 61, 98–114.

Dubbs, A., J. Guevara & R. Yuste (2016) moco: Fast Motion Correction for Calcium Imaging. Front Neuroinform, 10, 6.

Eisenhardt, M., A. C. Hansson, R. Spanagel & A. Bilbao (2015) Chronic intermittent ethanol exposure in mice leads to an up-regulation of CRH/CRHR1 signaling. Alcohol Clin Exp Res, 39, 752–62.

Fransson, S., A. Ruusala & P. Aspenstrom (2006) The atypical Rho GTPases Miro-1 and Miro-2 have essential roles in mitochondrial trafficking. Biochem Biophys Res Commun, 344, 500–10.

Goto, M., H. Kitamura, M. M. Alam, N. Ota, T. Haseba, T. Akimoto, A. Shimizu, T. Takano-Yamamoto, M. Yamamoto & H. Motohashi (2015) Alcohol dehydrogenase 3 contributes to the protection of liver from nonalcoholic steatohepatitis. Genes Cells, 20, 464–80.

Hollenbeck, P. J. (1996) The pattern and mechanism of mitochondrial transport in axons. Front Biosci, 1, d91–102.

Hollenbeck, P. J. & W. M. Saxton (2005) The axonal transport of mitochondria. J Cell Sci, 118, 5411–9.

Holtmaat, A., T. Bonhoeffer, D. K. Chow, J. Chuckowree, V. De Paola, S. B. Hofer, M. Hubener, T. Keck, G. Knott, W. C. Lee, R. Mostany, T. D. Mrsic-Flogel, E. Nedivi, C. Portera-Cailliau, K. Svoboda, J. T. Trachtenberg & L. Wilbrecht (2009) Long-term, high-resolution imaging in the mouse neocortex through a chronic cranial window. Nature protocols, 4, 1128–44.

Horishita, T. & R. A. Harris (2008) n-Alcohols inhibit voltage-gated Na+ channels expressed in Xenopus oocytes. J Pharmacol Exp Ther, 326, 270–7.

Iacobucci, G. J. & S. Gunawardena (2018) Ethanol stimulates the in vivo axonal movement of neuropeptide dense-core vesicles in Drosophila motor neurons. J Neurochem, 144, 466–482.

Jaiswal, A., W. J. Godinez, M. J. Lehmann & K. Rohr. 2016. Direct combination of multi-scale detection and multi-frame association for tracking of virus particles in microscopy image data. In Proc. of IEEE International Symposium on Biomedical Imaging (ISBI’16), 976–979. Prague, Czech Republic.

Kalman, R. E. (1960) A new approach to linear filtering and prediction problems. J. Basic Eng., D 82, 35–45.

Kang, J. S., J. H. Tian, P. Y. Pan, P. Zald, C. Li, C. Deng & Z. H. Sheng (2008) Docking of axonal mitochondria by syntaphilin controls their mobility and affects short-term facilitation. Cell, 132, 137–48.

Kiryu-Seo, S., N. Ohno, G. J. Kidd, H. Komuro & B. D. Trapp (2010) Demyelination increases axonal stationary mitochondrial size and the speed of axonal mitochondrial transport. J Neurosci, 30, 6658–66.

Knabbe, J., J. P. Nassal, M. Verhage & T. Kuner (2018) Secretory vesicle trafficking in awake and anaesthetized mice: differential speeds in axons versus synapses. J Physiol, 596, 3759–3773.

Kuner, T., R. Schoepfer & E. R. Korpi (1993) Ethanol inhibits glutamate-induced currents in heteromeric NMDA receptor subtypes. Neuroreport, 5, 297–300.

Kwinter, D. M., K. Lo, P. Mafi & M. A. Silverman (2009) Dynactin regulates bidirectional transport of dense-core vesicles in the axon and dendrites of cultured hippocampal neurons. Neuroscience, 162, 1001–10.

Laduron, P. M. & P. A. De Witte (1987) Enhanced axonal transport of receptor-bound opiate in ethanol-treated rats. Neurosci Lett, 77, 344–8.

Lamprecht, R. & J. LeDoux (2004) Structural plasticity and memory. Nat Rev Neurosci, 5, 45–54.

Lewis, T. L., Jr., G. F. Turi, S. K. Kwon, A. Losonczy & F. Polleux (2016) Progressive Decrease of Mitochondrial Motility during Maturation of Cortical Axons In Vitro and In Vivo. Curr Biol, 26, 2602–2608.

Liu, Y., G. Chen, C. Ma, K. A. Bower, M. Xu, Z. Fan, X. Shi, Z. J. Ke & J. Luo (2009) Overexpression of glycogen synthase kinase 3beta sensitizes neuronal cells to ethanol toxicity. J Neurosci Res, 87, 2793–802.

Llorens-Martin, M., G. Lopez-Domenech, E. Soriano & J. Avila (2011) GSK3beta is involved in the relief of mitochondria pausing in a Tau-dependent manner. PLoS One, 6, e27686.

Lovinger, D. M. & M. Roberto (2013) Synaptic effects induced by alcohol. Curr Top Behav Neurosci, 13, 31–86.

Lovinger, D. M., G. White & F. F. Weight (1989) Ethanol inhibits NMDA-activated ion current in hippocampal neurons. Science, 243, 1721–4.

Macaskill, A. F., J. E. Rinholm, A. E. Twelvetrees, I. L. Arancibia-Carcamo, J. Muir, A. Fransson, P. Aspenstrom, D. Attwell & J. T. Kittler (2009) Miro1 is a calcium sensor for glutamate receptor-dependent localization of mitochondria at synapses. Neuron, 61, 541–55.

Malatova, Z. & D. Cizkova (2002) Effect of ethanol on axonal transport of cholinergic enzymes in rat sciatic nerve. Alcohol, 26, 115–20.

Marszalec, W., Y. Kurata, B. J. Hamilton, D. B. Carter & T. Narahashi (1994) Selective effects of alcohols on gamma-aminobutyric acid A receptor subunits expressed in human embryonic kidney cells. J Pharmacol Exp Ther, 269, 157–63.

McCarron, J. G., C. Wilson, M. E. Sandison, M. L. Olson, J. M. Girkin, C. Saunter & S. Chalmers (2013) From structure to function: mitochondrial morphology, motion and shaping in vascular smooth muscle. J Vasc Res, 50, 357–71.

McLane, J. A. (1990) Retrograde axonal transport in chronic ethanol-fed and thiamine-deficient rats. Alcohol, 7, 103–6.

McLane, J. A., M. B. Atkinson, J. McNulty & A. C. Breuer (1992) Direct measurement of fast axonal organelle transport in chronic ethanol-fed rats. Alcohol Clin Exp Res, 16, 30–7.

Misgeld, T., M. Kerschensteiner, F. M. Bareyre, R. W. Burgess & J. W. Lichtman (2007) Imaging axonal transport of mitochondria in vivo. Nature methods, 4, 559–61.

Qiao, Q., L. Ma, W. Li, J. W. Tsai, G. Yang & W. B. Gan (2016) Long-term stability of axonal boutons in the mouse barrel cortex. Dev Neurobiol, 76, 252–61.

Rabin, R. A. & P. B. Molinoff (1981) Activation of adenylate cyclase by ethanol in mouse striatal tissue. J Pharmacol Exp Ther, 216, 129–34.

Rich, P. R. & A. Marechal (2010) The mitochondrial respiratory chain. Essays Biochem, 47, 1–23.

Sage, D., F. R. Neumann, F. Hediger, S. M. Gasser & M. Unser (2005) Automatic tracking of individual fluorescence particles: application to the study of chromosome dynamics. IEEE Trans Image Process, 14, 1372–83.

Schwenger, D. B. & T. Kuner (2010) Acute genetic perturbation of exocyst function in the rat calyx of Held impedes structural maturation, but spares synaptic transmission. Eur J Neurosci, 32, 974–84.

Smit-Rigter, L., R. Rajendran, C. A. Silva, L. Spierenburg, F. Groeneweg, E. M. Ruimschotel, D. Van Versendaal, C. van der Togt, U. T. Eysel, J. A. Heimel, C. Lohmann & C. N. Levelt (2016) Mitochondrial Dynamics in Visual Cortex Are Limited In Vivo and Not Affected by Axonal Structural Plasticity. Curr Biol, 26, 2609–2616.

Smith, H. L., J. N. Bourne, G. Cao, M. A. Chirillo, L. E. Ostroff, D. J. Watson & K. M. Harris (2016) Mitochondrial support of persistent presynaptic vesicle mobilization with age-dependent synaptic growth after LTP. Elife, 5.

Soderpalm, B., H. H. Lido & M. Ericson (2017) The Glycine Receptor-A Functionally Important Primary Brain Target of Ethanol. Alcohol Clin Exp Res, 41, 1816–1830.

Takihara, Y., M. Inatani, K. Eto, T. Inoue, A. Kreymerman, S. Miyake, S. Ueno, M. Nagaya, A. Nakanishi, K. Iwao, Y. Takamura, H. Sakamoto, K. Satoh, M. Kondo, T. Sakamoto, J. L. Goldberg, J. Nabekura & H. Tanihara (2015) In vivo imaging of axonal transport of mitochondria in the diseased and aged mammalian CNS. Proc Natl Acad Sci U S A, 112, 10515–20.

Verstreken, P., C. V. Ly, K. J. Venken, T. W. Koh, Y. Zhou & H. J. Bellen (2005) Synaptic mitochondria are critical for mobilization of reserve pool vesicles at Drosophila neuromuscular junctions. Neuron, 47, 365–78.

Yi, M., D. Weaver & G. Hajnoczky (2004) Control of mitochondrial motility and distribution by the calcium signal: a homeostatic circuit. J Cell Biol, 167, 661–72.

Zahn, T. R., J. K. Angleson, M. A. MacMorris, E. Domke, J. F. Hutton, C. Schwartz & J. C. Hutton (2004) Dense core vesicle dynamics in Caenorhabditis elegans neurons and the role of kinesin UNC-104. Traffic, 5, 544–59.

Zorumski, C. F., S. Mennerick & Y. Izumi (2014) Acute and chronic effects of ethanol on learning-related synaptic plasticity. Alcohol, 48, 1–17.

